# In vivo discovery of blood-brain barrier opening small molecules with FishNAP

**DOI:** 10.64898/2026.03.18.712473

**Authors:** Thomas C. Potts, Erin E. McDonnell, Jacklyn Levey, Lewis G. Gall, Evangeline Coffinas, Colleen G. Rutley, Neha Gururaj, Alexandra K. Vindigni, Advait R. Iyer, Kaitlyn A. Pericone, Manan H. Gosai, Megan E. Canace, Natasha M. O’Brown

## Abstract

The blood-brain barrier (BBB) maintains neural homeostasis by tightly regulating molecular exchange between the circulation and the brain, but it also restricts drug delivery and thereby limits treatment of neurological disorders. Pharmacological agents that transiently and safely increase BBB permeability could therefore improve brain drug delivery, yet their systematic discovery has been constrained by the limitations of existing approaches. Here we present FishNAP, a non-invasive, high-throughput zebrafish platform for real-time *in vivo* assessment of BBB permeability. Screening 2,320 FDA-approved small molecules identified seven that increased BBB permeability in zebrafish, with barrier integrity recovering within 24 hours in each case, indicating reversible modulation. Three of these compounds, Calcitriol, Lovastatin and Sunitinib, which act through distinct signaling pathways, also increased BBB permeability in adult mice through both tight junction disruption and increased transcytosis. FishNAP thus enables the identification of conserved modulators of BBB function with translational potential for brain drug delivery.

---

The blood–brain barrier (BBB) arises from specialized properties of the brain vasculature that tightly regulate molecular and cellular exchange between the circulation and brain parenchyma. This selective permeability is essential for maintaining the brain’s homeostatic environment and protecting neuronal tissue from toxins and pathogens^1^. Barrier function arises from tight junctions between endothelial cells, suppression of vesicular transcytosis, and specialized transport systems that regulate nutrient influx and waste removal^2^. While these features are critical for neural health, they also present a major obstacle for central nervous system (CNS) therapeutics—over 98% of small molecules fail to reach the brain at pharmacologically relevant concentrations^3^. Consequently, substantial effort has focused on strategies to bypass or transiently modulate the BBB, including chemical modification of drugs to enhance penetration, receptor-mediated transport, and focused ultrasound to transiently disrupt tight junctions^4,5^. Although these approaches have shown promise in specific contexts, identifying compounds capable of safely and reversibly modulating the intact BBB remains challenging.

A number of experimental platforms have been developed to study BBB permeability and evaluate candidate therapeutics. *In vitro* systems—including transwell assays, organ-on-chip technologies, and human induced pluripotent stem cell (iPSC)-derived neurovascular units—have provided valuable insight into endothelial barrier biology and offer tractable platforms for mechanistic studies and drug testing in the human context^6,7^. However, these models do not fully capture the complex cellular interactions and physiological forces present in the intact brain vasculature. Indeed, the maintenance of BBB identity depends on signaling between endothelial cells and surrounding cells in the brain, including pericytes, neurons, and astrocytes^8–12^. These interactions are difficult to fully reproduce *in vitro*, resulting in a rapid loss of these BBB molecular signatures^13^. As a result, there remains a need for experimental systems that preserve the physiological context of the BBB while remaining compatible with scalable discovery approaches.

Zebrafish provide an opportunity to bridge this gap. The zebrafish BBB shares key molecular and structural features with the mammalian barrier and becomes functionally restrictive by 5 days post-fertilization (dpf)^1,14,15^. In addition to this evolutionary conservation, zebrafish are highly tractable genetic and chemical systems: their external development, small size, and capacity for direct compound absorption from the surrounding water enable rapid *in vivo* manipulation and screening^16^. However, most existing zebrafish assays for BBB permeability rely on tracer injections or confocal imaging^8,15,17^, which substantially limit experimental throughput.

To address this limitation, we developed FishNAP, a zebrafish-based platform for high-throughput measurement of BBB permeability *in vivo*. FishNAP leverages the small size of zebrafish larvae and the rapid functional maturation of their BBB by 5 dpf^15^ to measure barrier permeability in a 96-well format using a behavioral phenotype that correlates with BBB leakage. Because this assay does not require tracer injections or high-resolution imaging, it enables rapid and scalable assessment of BBB modulation across large chemical libraries. Using FishNAP, we screened 2,320 FDA-approved small molecules and identified 11 compounds that reproducibly increased BBB permeability. Follow-up tracer leakage assays narrowed this to 7 molecules that robustly opened the zebrafish BBB. Importantly, three compounds (Calcitriol, Lovastatin, and Sunitinib), representing distinct signaling pathways, also increased BBB permeability in adult mice, indicating conservation of barrier modulation across vertebrates. Together, these findings establish FishNAP as a scalable *in vivo* platform for discovering pharmacological modulators of BBB permeability and identify candidate compounds capable of transiently opening the intact barrier.

## RESULTS

### Novel FishNAP assay for BBB function

To enable medium-throughput, *in vivo* assessment of BBB function, we developed a scalable behavioral screening platform called FishNAP (**Fish**-based assay for **N**on-invasive **A**ssessment of **P**ermeability). This assay uses larval zebrafish and measures BBB integrity based on loperamide-induced sedation in a 96-well format using Daniovision (Fig. 1A). Loperamide is a synthetic opioid receptor agonist that does not cross the intact BBB. Thus, larvae with a sealed BBB remain behaviorally active after exposure, whereas larvae with a leaky BBB exhibit reduced activity as loperamide enters the brain and induces sedation. We first validated FishNAP across developmental stages with distinct BBB states. At 3 and 4 dpf, when elevated endothelial transcytosis renders the BBB functionally immature^15^, larvae exhibited robust loperamide-induced sedation (Fig. 1B). In contrast, at 5-7 dpf, when transcytosis is suppressed and the BBB has functionally sealed^15^, larvae remained behaviorally active following loperamide exposure (Fig. 1B), consistent with an intact barrier. Importantly, larvae at all stages remained equally active in the absence of loperamide (Fig. S1). Interestingly, larvae at 6 dpf displayed a significantly higher activity after exposure to loperamide than at 5 and 7 dpf (Fig. 1B), which may reflect developmental refinement of inhibitory motor circuits by 7 dpf^18^. To further validate the assay, we tested the zebrafish *spock1* mutant line, which has regionally restricted BBB permeability in the forebrain and midbrain due to defective tight junctions and elevated transcytosis^8^, and found that, compared to wild-type controls, *spock1* mutants displayed loperamide-induced sedation (Fig. 1C), demonstrating that FishNAP sensitively detects both developmental and genetically induced BBB dysfunction.

**Figure 1.**
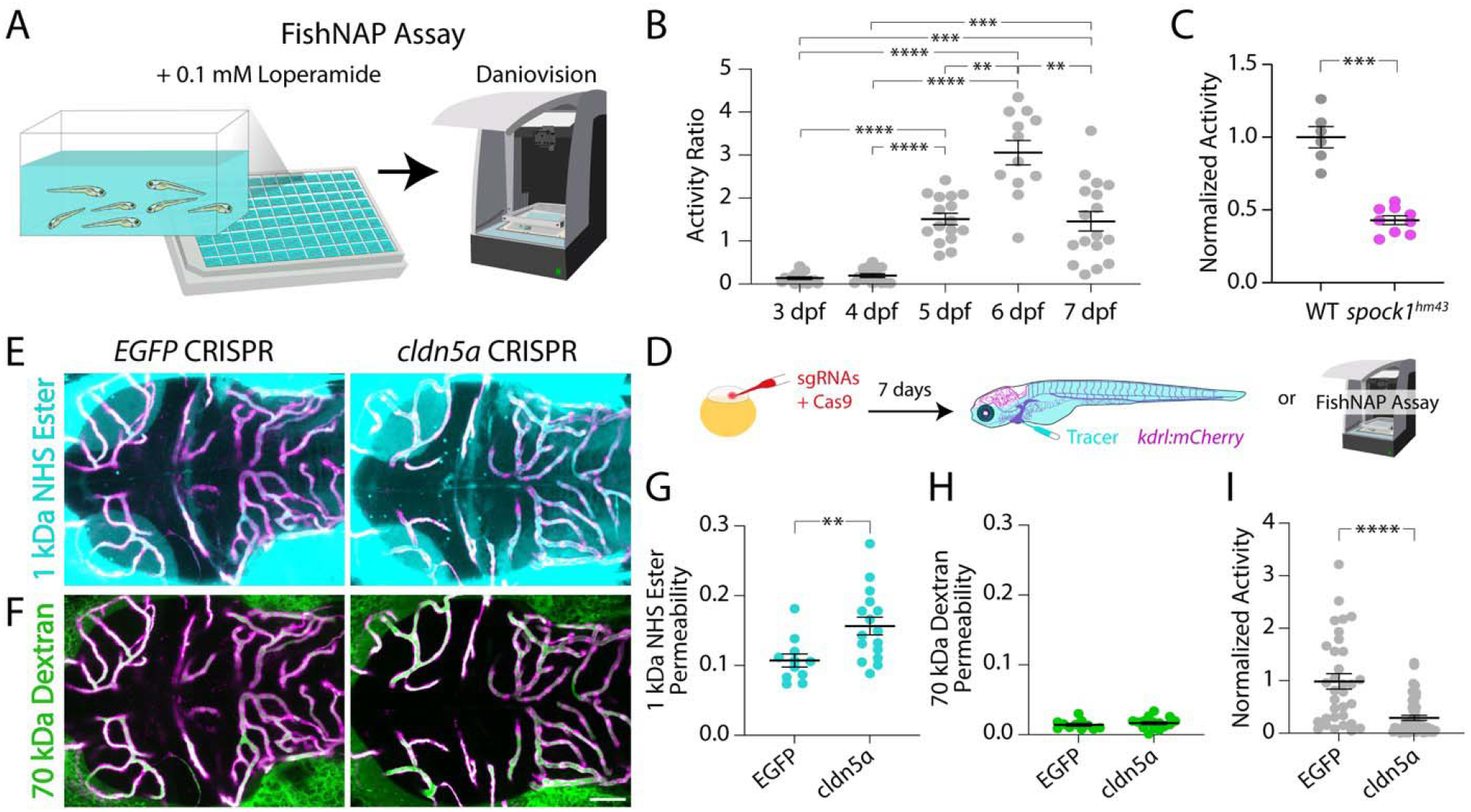
FishNAP reliably reveals increased BBB permeability. **A**.Schematic of the FishNAP assay, with up to seven larvae per well. Overall activity is monitored using Daniovision for 1 hour after loperamide exposure; increased BBB permeability leads to reduced locomotor activity. **B**. Leaky 3 and 4 dpf larvae are significantly sedated compared to 5-7 dpf fish, as previously reported^15^. **C**. *spock1* mutants (magenta) show significantly reduced activity compared to wild-type (WT) controls at 7 dpf using FishNAP. **D**. Workflow schematic for assessing BBB permeability in *cldn5a* crispant fish. **E–H**. Tracer leakage assays using 1 kDa NHS Ester (E, turquoise) and 70 kDa Dextran (F, green) reveal significantly increased 1 kDa NHS tracer permeability outside of the *kdrl:mCherry*+ vasculature (magenta) in *cldn5a* crispants (N=16) vs. *EGFP* crispant controls (N=11, G), but no significant change in 70 kDa Dextran permeability (H). **I**. FishNAP also detects increased BBB permeability in *cldn5a* crispants (N=60) relative to *EGFP* crispant controls (N=36). Circle data points represent a single well containing 7 fish, with a minimum of 6 wells (42 larvae) per treatment (B and C) and square data points represent individual larvae (G-I). Bars show the mean ± SEM; ** p<0.01, *** p<0.001, **** p<0.0001 by Welch ANOVA with Dunnett’s T3 post-hoc test (B), two-sided Welch’s t-test (C, G) or two-sided Mann-Whitney test (I). Scale bar represents 50 µm. Source data are provided in a Source Data file.

We further tested the FishNAP assay on *cldn5a* F0 crispants, which exhibit mosaic disruption of the tight junction gene *cldn5a* that is essential for BBB integrity^19^. BBB function in these fish was assessed using both functional tracer leakage assays and FishNAP (Fig. 1D). Permeability to a small 1 kDa NHS Ester tracer (Fig. 1E) and a larger 70 kDa Dextran tracer (Fig. 1F) revealed a significant increase in 1 kDa NHS Ester tracer permeability (Fig. 1G), but no significant change in 70 kDa Dextran permeability in the *cldn5a* crispants compared to *EGFP* crispant controls (Fig. 1H), consistent with altered tight junction function in the mosaic *cldn5a* crispants, which had an average of 60% knockdown of *cldn5a* expression compared to controls (Fig. S2). Traditional fluorescent tracer leakage assays required approximately three hours for injection, circulation, mounting, and confocal imaging (Fig. 1E–H), whereas FishNAP detected significant BBB impairment in more *cldn5a* crispants compared to control-injected siblings within just one hour (Fig. 1I). To assess whether *cldn5a* knockdown corresponded to lower activity ratios, we performed qPCR on individual larvae following FishNAP screening; however, we did not observe a significant correlation between *cldn5a* transcript levels and activity ratio (Fig. S2). Even without this correlation, FishNAP could still rapidly distinguish larvae with significant BBB phenotypes among F0 crispants (Fig. 1I). Because this assay is nonlethal and does not require phenylthiourea (PTU) treatment, which is typically used to block pigmentation for confocal imaging in wild-type fish, it enables rapid and scalable screening of BBB function. Notably, larvae showing strong BBB phenotypes could be placed directly on the fish system, streamlining the generation of stable mutant lines by allowing early selection and rearing of the most penetrant individuals.

### Identification of BBB-opening small molecules

Using the FishNAP platform, we screened a library of 2,320 FDA-approved small molecules (ApexBio, Supplementary Data 1), to identify compounds capable of opening a sealed BBB. Zebrafish were treated in bath with two pooled compounds, each at 5 μM, from 5 to 7 dpf, and BBB integrity was assessed using FishNAP (Fig. 2A). Putative hits were defined as having a normalized activity ratio below 0.6. Following pooled testing, we deconvoluted the pools by individually testing compounds from toxic or hit pairs. This yielded 36 molecules that induced sedation alone, as well as 420 molecules that remained toxic at 5 μM (Fig. 2B). These 420 toxic molecules were then retested individually at 1 μM, which revealed an additional 58 compounds capable of inducing sedation at the lower dose and 37 compounds that were still toxic at 1 μM. Compounds intended solely for topical use, antiseptics, or food additives were excluded from further testing at this stage. The remaining toxic molecules were retested at 0.5 μM, which revealed another 7 small molecules that reduced activity below our 0.6 threshold. All 101 putative hits were then reviewed for reasonable potential as a systemic treatment in humans before selection for follow-up testing and hit confirmation. The remaining candidates were tested in at least three biological replicates to confirm reproducibility, resulting in 11 small molecules that consistently caused BBB leakage in the FishNAP assay after 48 hours of treatment (Fig. 2C). To assess the kinetics of BBB disruption, we tested these 11 molecules after just 24 hours of exposure. We found that 10 of the 11 compounds (all except for Afatinib dimaleate) induced sedation within the shorter time window (Fig. 2D). Together, these results demonstrate the power of FishNAP for *in vivo* screening of BBB-modulating small molecules.

**Figure 2.**
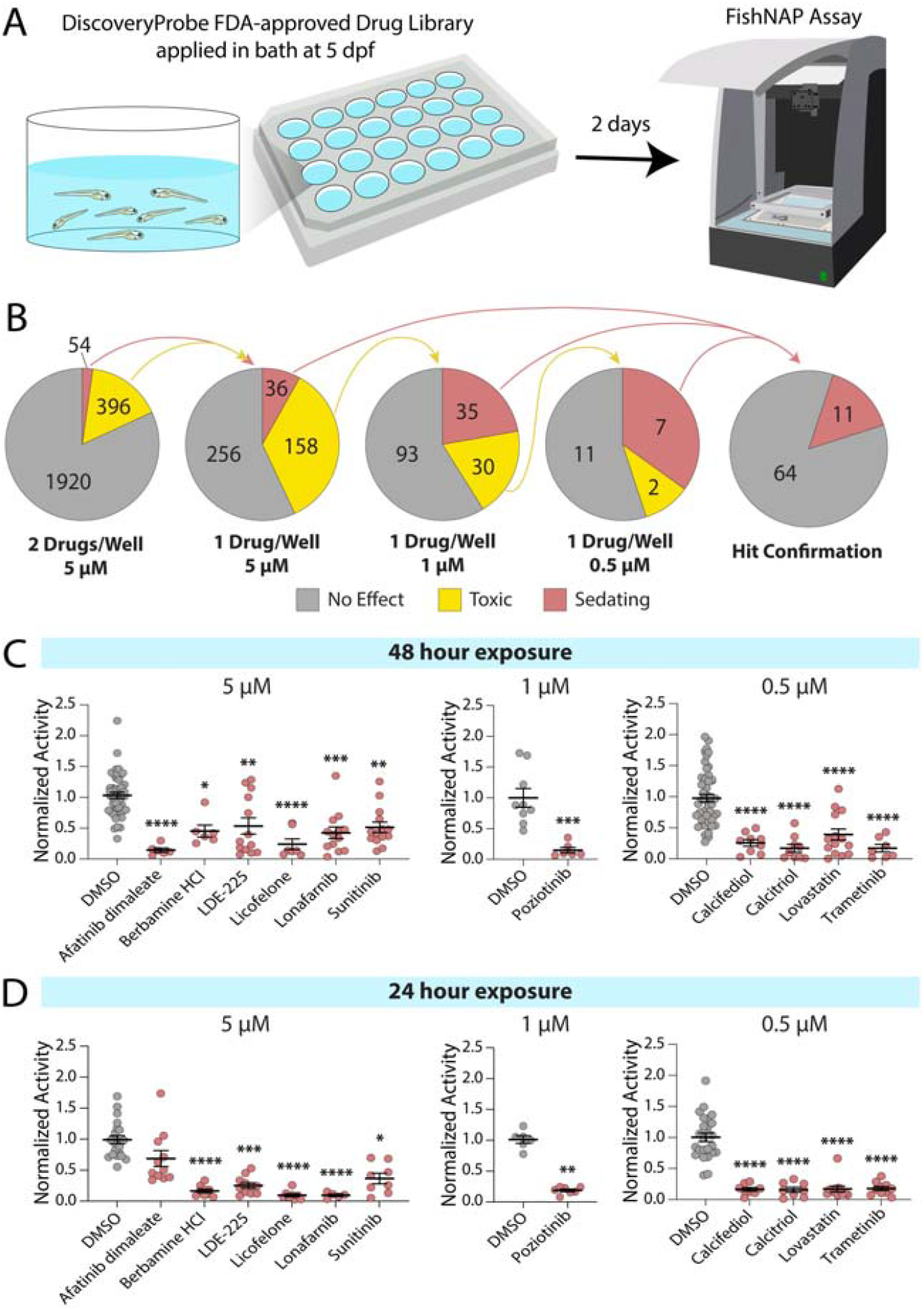
FishNAP-based screening of FDA-approved small molecules identifies 11 compounds that significantly reduce locomotor activity in zebrafish larvae. **A**.Schematic of the small-molecule treatment paradigm in which seven larvae were treated from 5 to 7 dpf in a 24-well plate and assayed for BBB function using FishNAP at 7 dpf. **B**. Pie charts illustrating the screening workflow, beginning with pooled testing at 5 µM, followed by individual compound testing at 5 µM and lower concentrations (1 µM and 0.5 µM) for compounds that were toxic (yellow). Putative hits (puce) were validated by replicate testing, yielding 11 confirmed small-molecule hits. **C**. Validation of the 11 compounds that significantly sedated larvae after 48 hours of treatment compared to DMSO controls at their respective concentrations. **D**. FishNAP reveals that 10 of these compounds can induce sedation even after only 24 hours of treatment. Each point represents an individual well of 7 fish, with a minimum sample size of 6 wells (42 fish) for each treatment. Bars show the mean ± SEM; * p<0.05, ** p<0.01, *** p<0.001, **** p<0.0001 by Kruskal-Wallis test with Dunn’s post-hoc test to DMSO controls (5 µM and 0.5 µM) or Mann-Whitney test (1 µM). Source data are provided in a Source Data file.

To confirm that the 11 identified compounds caused sedation through increased BBB permeability rather than off-target effects, we repeated the 48-hour treatment paradigm followed by functional tracer leakage assays. We used two molecular weight tracers, 1 kDa NHS Ester and 10 kDa Dextran, to determine not only whether the BBB was compromised but also the likely subcellular mechanism, since tight junction defects selectively increase permeability to the 1 kDa tracer^20^, as seen in the *cldn5a* crispants (Fig. 1E-H) whereas elevated transcytosis increases permeability to both^15^. These assays revealed that 8 of the 11 compounds significantly increased BBB permeability compared with DMSO-treated controls (Fig. 3, Fig. S3), and seven—Calcifediol, Calcitriol, LDE-225, Lonafarnib, Lovastatin, Sunitinib, and Trametinib—caused substantial increases in 1 kDa NHS Ester permeability (Fig. 3A-B), and one, Licofelone, cause significant but small increases in 1 kDa tracer permeability (Fig. S3). Most of these compounds also enhanced permeability to the larger 10 kDa Dextran tracer, except for LDE-225 and Trametinib (Fig. 3C-D), which likely act exclusively through tight junction disruption rather than increased transcytosis. Calcitriol, Calcifediol, and Sunitinib produced the greatest increases in 10 kDa Dextran permeability, while Lovastatin and Lonafarnib induced smaller but significant effects. To assess whether these effects were specific to the BBB rather than reflecting a generalized increase in vascular permeability, we measured 1 kDa NHS Ester and 10 kDa Dextran leakage in two peripheral tissues, trunk fin and muscle. Because peripheral vasculature in fin and muscle is inherently more permeable than the BBB, subtle changes may be harder to detect against that baseline; nonetheless, none of the small molecules produced an overt increase in permeability in either tissue (Fig. S4), supporting an effect enriched at the BBB rather than broadly systemic.

**Figure 3.**
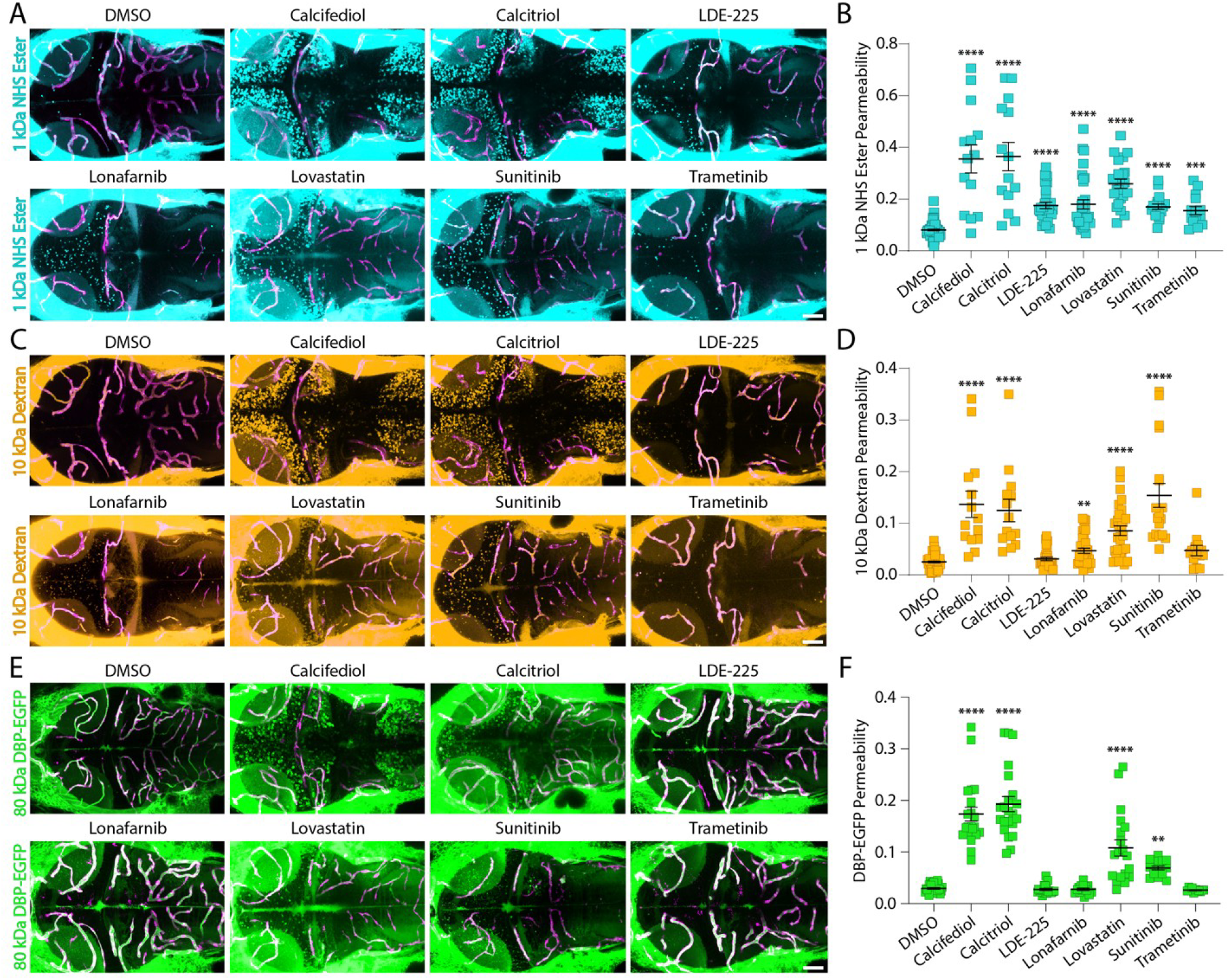
Functional tracer leakage assays validate small molecule opening of the BBB. **A-B**.Larvae were treated with each small molecule from 5–7 dpf and assayed for leakage of 1 kDa NHS Ester (turquoise) from the vasculature (magenta) into the brain (A); all seven compounds significantly increased permeability to this small tracer (B). **C-D**. The same treatment paradigm was used to assess leakage of 10 kDa Dextran (gold) from the vasculature into the brain (C), revealing increased permeability relative to DMSO controls for all compounds except LDE-225 and Trametinib (D). **E-F**. Treated larvae were also assessed for leakage of the large, endogenous 80 kDa DBP-EGFP tracer (green) from the vasculature (E), revealing a significant increase in Calcifediol, Calcitriol, Lovastatin, and Sunitinib treated fish (F). Each data point represents an individual fish, with a minimum of 14 larvae per treatment. Bars show the mean ± SEM; * p<0.05, ** p<0.01, *** p<0.001, **** p<0.0001 by Kruskal-Wallis test with Dunn’s post-hoc test compared to DMSO controls. Scale bars represent 50 µm. Source data are provided in a Source Data file.

To further resolve the extent of barrier disruption, we extended the tracer panel for these seven compounds to include a smaller 376 Da fluorescein sodium tracer and a larger 80 kDa endogenous DBP-EGFP tracer. All seven compounds increased permeability to fluorescein (Fig. S5), consistent with paracellular disruption extending to even smaller size tracers than the 1 kDa NHS (Fig. 3A). By contrast, Calcifediol, Calcitriol, and Lovastatin produced substantial increases in DBP-EGFP permeability (Fig. 3E-F), indicating a greater degree of barrier disruption sufficient to admit this larger endogenous tracer, and Sunitinib produced a smaller but significant increase in signal (Fig. 3F). These hits targeted diverse signaling pathways, including Hedgehog (LDE-225), farnesyltransferase/Ras (Lonafarnib), HMG-CoA reductase/cholesterol (Lovastatin), multi-kinase RTK/VEGF (Sunitinib), and MEK/ERK MAPK (Trametinib), with vitamin D signaling (Calcifediol and Calcitriol) as the only pathway with multiple hits. Notably, both vitamin D agonists also increased BBB permeability throughout the brain, implicating vitamin D in BBB opening for the first time. To determine whether these effects extended to other members of the same signaling pathways, we screened 51 additional compounds from these pathways that had not met the original hit criteria and identified 10 more putative BBB regulators: three additional Hedgehog/Smoothened inhibitors, one additional farnesyltransferase inhibitor (Tipifarnib), five additional HMG-CoA reductase inhibitors, one additional MEK inhibitor, and one additional VEGFR inhibitor (Fig. S6). However, this activity was not shared uniformly across all pathway members, with the exception of the farnesyltransferase inhibitors, where Tipifarnib also produced an effect similar to Lonafarnib. This suggests that BBB opening is not necessarily a pathway-specific effect but may instead depend on compound-specific properties within a given pathway.

### Recovery of BBB integrity after drug-induced opening

To assess whether the effects of these seven BBB-modulating small molecules were reversible, we treated larvae with each compound for 24 hours, which was sufficient to increase BBB permeability (Fig. 2D), then washed into fresh, drug-free media and monitored daily using FishNAP to determine when BBB integrity was restored (Fig. 4A). This revealed full recovery within 24 hours for Calcifediol, LDE-225, Lonafarnib, Sunitinib, and Trametinib (Fig. 4B). However, Calcitriol and Lovastatin at the screening concentration of 0.5 µM failed to recover within 48 hours (Fig. 4B), suggesting either irreversible BBB disruption or prolonged drug retention that maintains pharmacological activity during washout. We therefore determined minimal effective doses to open the BBB within 24 hours using FishNAP for all seven compounds (Fig. 4C-D, Fig. S7). This revealed that Calcifediol, Lonafarnib, and Trametinib remained effective at 0.05 µM, LDE-225 was effective only at the screening concentration of 0.5 µM, and Sunitinib required substantially higher concentrations, remaining effective at 2.5 µM (Fig. S7). For Calcitriol and Lovastatin, which did not fully recover at the screening concentration, we observed significant effects at both 0.05 and 0.005 µM for Calcitriol (Fig. 4C) and at 0.25 µM for Lovastatin (Fig. 4D). Tracer leakage assays validated significant permeability at 0.05 µM Calcitriol, but not 0.005 µM (Fig. 4E-F), indicating FishNAP’s greater sensitivity to subtle BBB changes. We also observed significantly increased permeability at 0.25 µM and 0.5 µM Lovastatin within 24 hours of treatment (Fig. 4E-F). Critically, repeating reversibility assays at 0.05 µM Calcitriol and 0.25 µM Lovastatin showed full recovery within 24 hours (Fig. 4G), confirming that all seven compounds reversibly modulate the BBB when administered at doses that minimize irreversible barrier damage.

**Figure 4.**
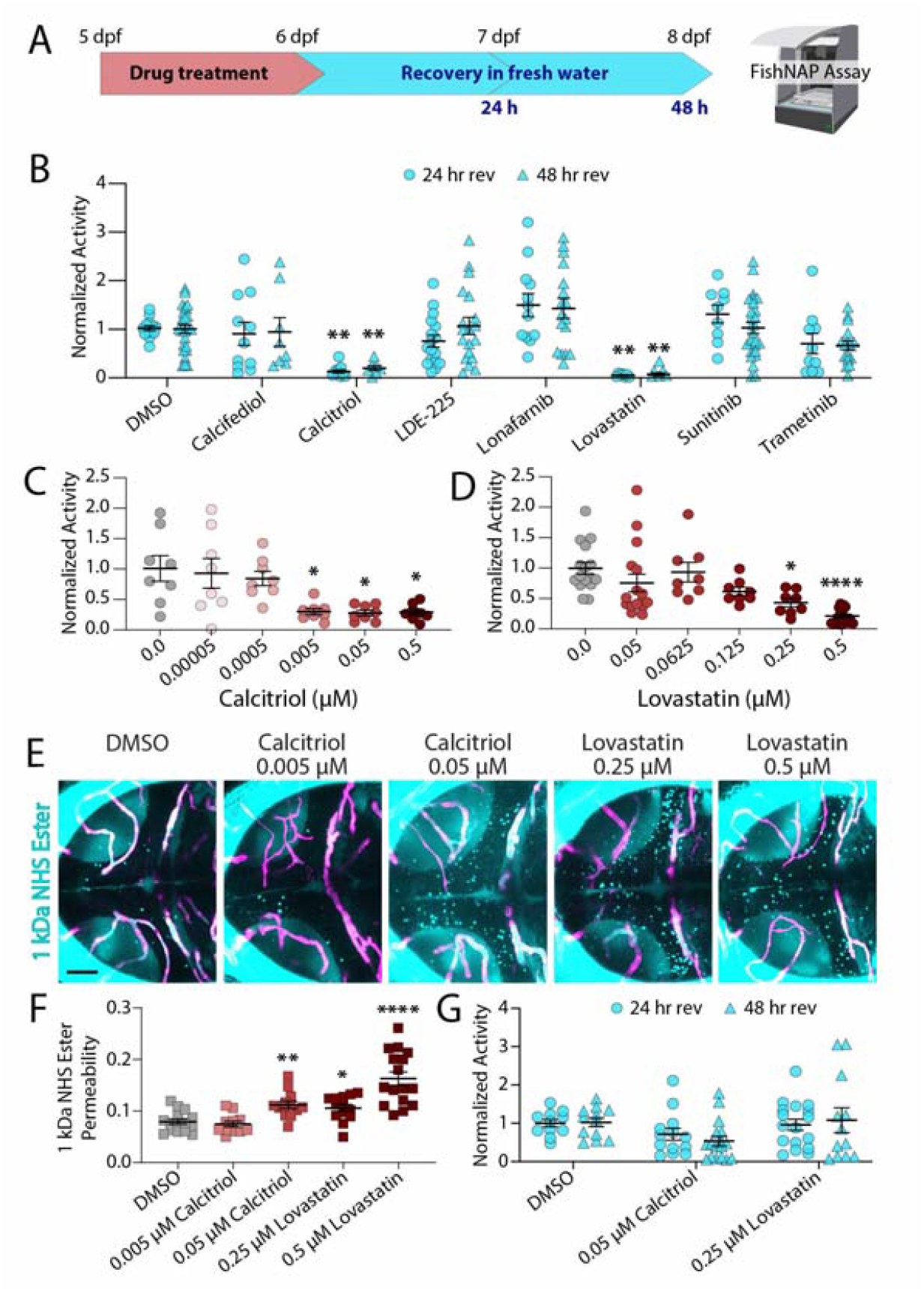
BBB modulation by these seven small molecules is transient. **A**.Schematic of small molecule treatment and recovery paradigm after 24 hours of small molecule exposure. **B**. Recovery assays reveal that five of the BBB-opening small molecules are able to recover full BBB function within 24 hours (Calcifediol, LDE-225, Lonafarnib, Sunitinib, and Trametinib) at the screening concentrations, while Calcitriol and Lovastatin remain leaky for 48 hours after 0.5 µM exposures. **C-D**. Dose-response curves for Calcitriol (C) and Lovastatin (D) reveal Calcitriol is able to significantly sedate at 0.005 µM and Lovastatin is able to sedate at 0.25 µM. **E-F**. Functional tracer leakage assays with 1 kDa NHS Ester (turquoise, E) reveal 0.05 µM Calcitriol exposure and 0.25 µM Lovastatin exposure are able to significantly increase BBB permeability outside of the vasculature (magenta), while 0.005 µM Calcitriol exposure does not have a significant effect (F). **G**. Recovery assays at lower doses of Calcitriol and Lovastatin reveal BBB functional recovery within 24 hours. Each data point represents an individual well of 7 fish (B-D, G) with a minimum sample size of 6 wells (42 fish) for each treatment or an individual fish (F), with a minimum of 10 larvae per condition. Bars show the mean ± SEM; * p<0.05, ** p<0.01, *** p<0.001, **** p<0.0001 by two-way mixed ANOVA with Dunnett’s post-hoc test compared to DMSO controls (B, G), Kruskal-Wallis test with Dunn’s post-hoc test compared to DMSO controls (C, D) or by Welch ANOVA with Dunnett’s T3 post-hoc test (F). Scale bar represents 50 µm. Source data are provided in a Source Data file.

### Assessing translational potential of BBB-opening small molecules in mammals

Finally, while our FishNAP assay enabled *in vivo* screening for BBB modulators in zebrafish (Fig. 2), it remained unclear whether these compounds would similarly affect the mammalian BBB. To test for cross-species conservation, we evaluated three of the BBB-opening small molecules identified by FishNAP that permeabilized the BBB to all tracers and acted through distinct signaling pathways—Calcitriol, Lovastatin, and Sunitinib—in adult mice. Mice were treated with each small molecule for two days prior to tracer leakage assays using the small 550 Da NHS-biotin tracer (Fig. 5A). While DMSO-treated controls confined the injected NHS tracer within their CD31+ vasculature (Fig. 5B), Calcitriol-, Lovastatin-, and Sunitinib-treated mice exhibited significant increases in tracer leakage (Fig. 5C–F). To determine whether this increased permeability extended to larger, physiologically relevant plasma proteins, we also assessed leakage of endogenous Albumin (66 kDa) and IgG (150 kDa). All three small molecules significantly increased Albumin accumulation in the brain, with Calcitriol producing the greatest leakage, Lovastatin a more moderate increase, and Sunitinib the least (Fig. 5G–K), mirroring the graded response we observed with the 80 kDa DBP-EGFP tracer in zebrafish (Fig. 3E-F). For the larger IgG molecule, however, only Calcitriol and Lovastatin significantly increased brain accumulation, again with Calcitriol producing more leakage than Lovastatin, while Sunitinib had no effect compared to DMSO controls (Fig. 5L–P).

**Figure 5.**
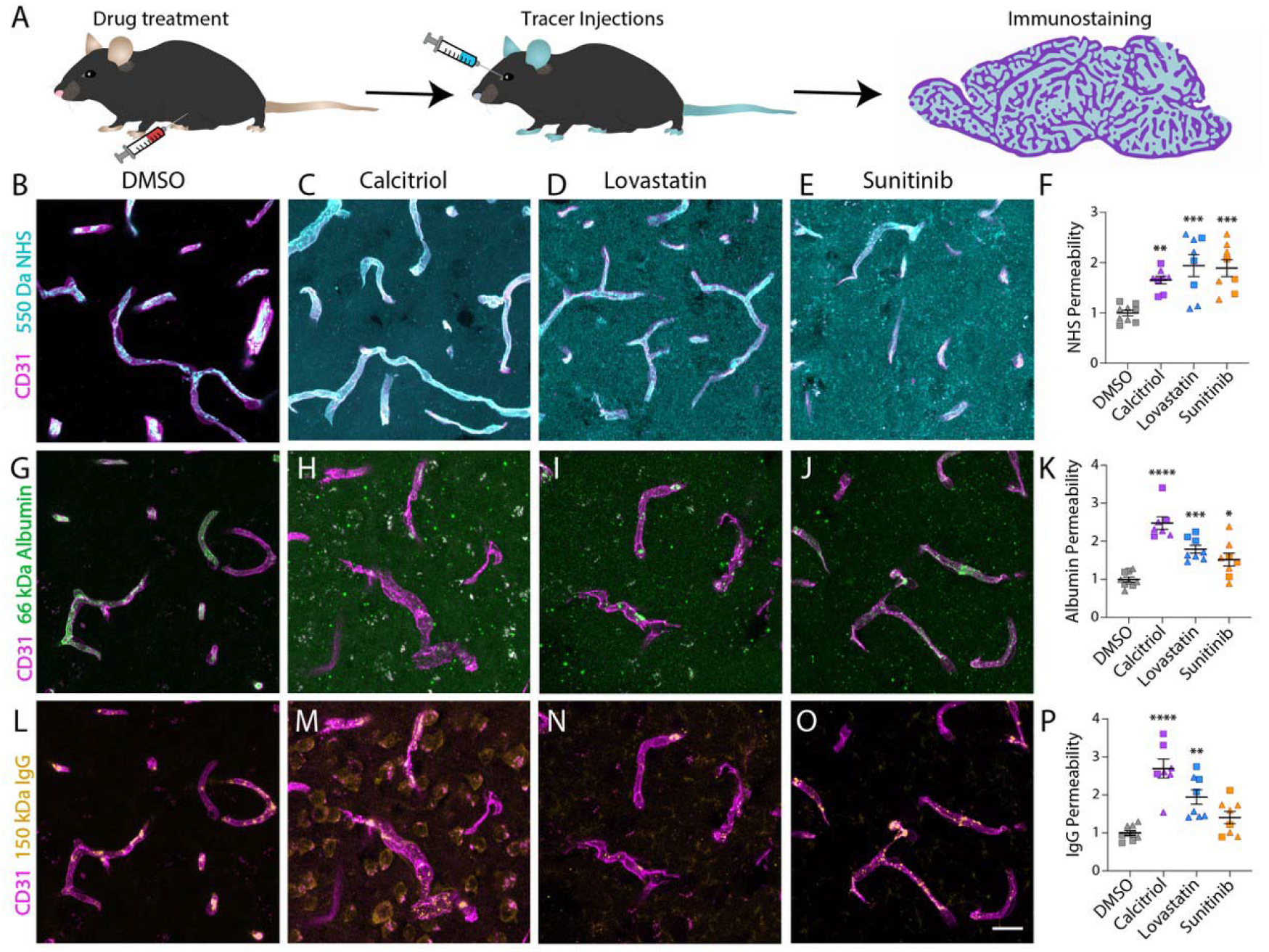
Small molecule treatment increases BBB permeability in adult mice. **A**.Schematic of drug treatment paradigm and downstream permeability assays. **B-F**. Tracer leakage assays using 550 Da NHS-Biotin (cyan) reveal a significant increase in tracer permeability outside the CD31+ vasculature (magenta) in Calcitriol (C), Lovastatin (D), and Sunitinib (E) treated mice relative to DMSO controls (B); quantified in F. **G-K**. Calcitriol (H), Lovastatin (I), and Sunitinib (J) increase extravasation of 66 kDa Albumin (green) into the brain parenchyma relative to DMSO controls (G), to varying degrees; quantified in K. **L-P**. Calcitriol (M) and Lovastatin (N), but not Sunitinib (O), increase extravasation of 150 kDa IgG (gold) relative to DMSO controls (L); quantified in P. Individual data points represent the average of three cortical area quantifications per mouse (minimum of 7 animals per condition), with males shown as triangles and females as squares in F, K, and P. Bars show the mean ± SEM; * p<0.05, ** p<0.01, *** p<0.001, **** p<0.0001 by one-way ANOVA with Dunnett’s post-hoc test to DMSO controls. Scale bar represents 20 µm. Source data are provided in a Source Data file.

Having established that these three small molecules increase BBB permeability in mice, we next sought to understand the cellular mechanisms underlying this effect. To determine whether this enhanced BBB permeability arose from tight junction dysfunction, increased transcytosis, or both, we immunostained treated mouse cortex for the tight junction protein CLDN5, the transcytotic suppressor MFSD2A, and the caveolar structural protein CAV1 (Fig. 6). Calcitriol-, Lovastatin-, and Sunitinib-treated mice exhibited significant decreases in endothelial CLDN5 and MFSD2A expression, along with a corresponding increase in CAV1 expression (Fig. 6), demonstrating that these three compounds disrupt the mammalian BBB through both tight junction breakdown and increased transcytosis even though they are in different signaling pathways, and that the mechanisms uncovered in zebrafish can translate directly to the mammalian BBB.

**Figure 6.**
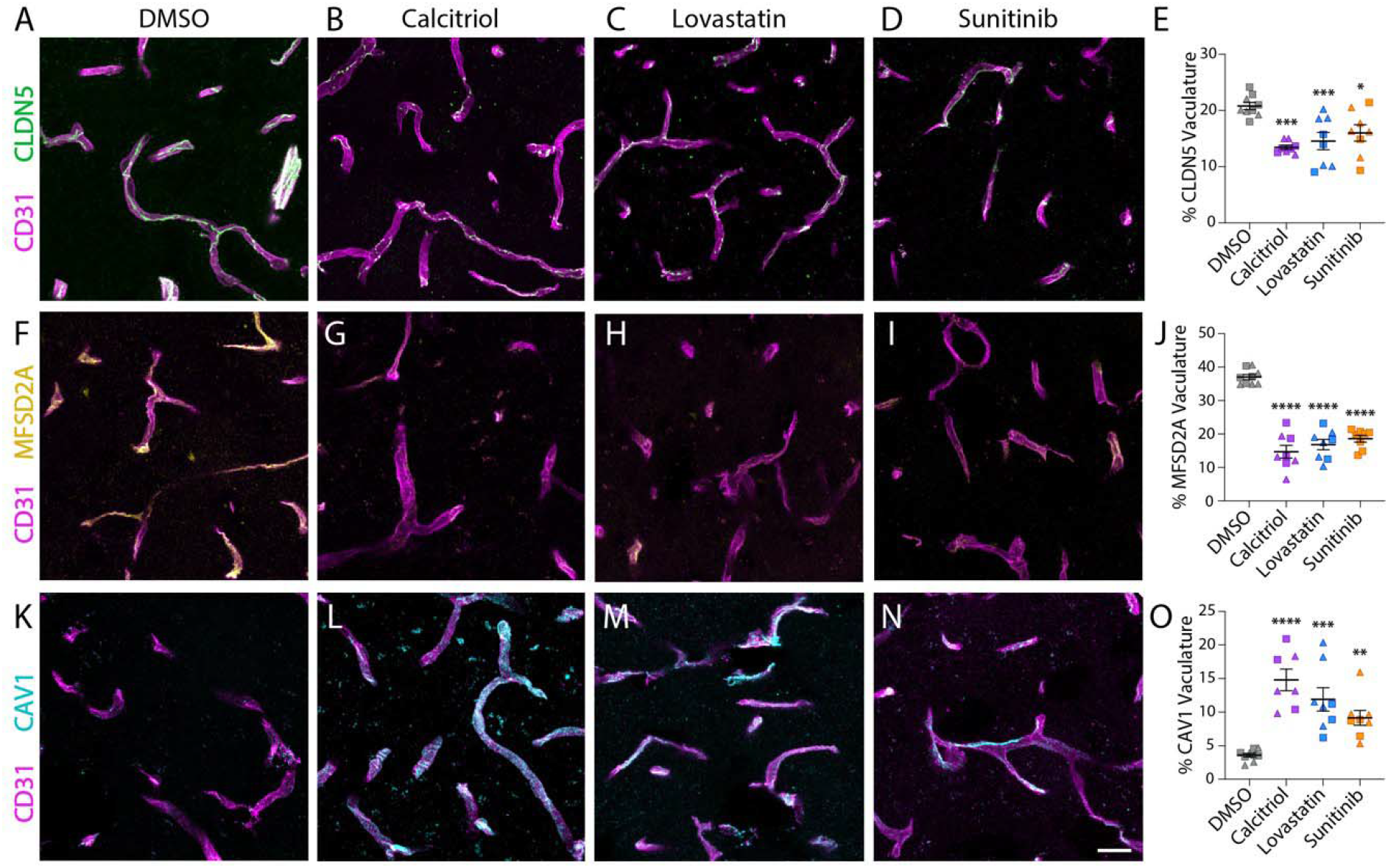
Small molecule treatment increases mouse BBB permeability through both tight junction breakdown and increased transcytosis. **A-E.**Calcitriol (B), Lovastatin (C), and Sunitinib (D) reduced endothelial (CD31, magenta) expression of the tight junction protein CLDN5 (green) relative to DMSO controls (A), which maintained a continuous, string-like pattern; quantified in E. **F-J**. The same treatments reduced endothelial expression of the transcytosis suppressor MFSD2A (gold) (G-I) relative to DMSO controls (F), which maintained MFSD2A expression throughout the cortical vasculature (magenta); quantified in J. **K-O**. Calcitriol (L), Lovastatin (M), and Sunitinib (N) significantly increased endothelial expression of the caveolae-forming protein CAV1 (cyan) relative to DMSO controls (K), which maintained low CAV1 expression; quantified in O. Individual data points represent the average of three cortical area quantifications per mouse (minimum of 7 animals per condition), with males shown as triangles and females as squares in E, J, and O. Bars show the mean ± SEM; * p<0.05, ** p<0.01, *** p<0.001, **** p<0.0001 by one-way ANOVA with Dunnett’s post-hoc test to DMSO controls. Scale bar represents 20 µm. Source data are provided in a Source Data file.

## DISCUSSION

While zebrafish are a powerful model for studying BBB development and function^8,14,15,17,21^, most existing assays rely on invasive tracer injections or imaging-based approaches that are difficult to scale. Although these methods have yielded important insights, they are not well-suited for large-scale genetic or small-molecule screens. This is unfortunate, as zebrafish larvae readily absorb compounds directly from the water, making them highly tractable for *in vivo* pharmacology and chemical screening^16,22^. One persistent challenge in BBB drug development is the gap between *in vitro* activity and *in vivo* efficacy^23^. While many compounds modulate BBB permeability in *in vitro* models, they frequently fail to penetrate the barrier *in vivo*^24^. *In vitro* systems are amenable to high-throughput screening but lack the cellular interactions, vascular architecture, hemodynamic forces, and whole-organism pharmacokinetics that shape BBB function. FishNAP addresses this gap by enabling rapid, nonlethal measurement of BBB permeability through a behavioral assay performed in a 96-well format in under an hour, without microinjection, confocal imaging, or extensive image analysis.

Using FishNAP, we screened a 2,320 compound FDA-approved library in under three months and identified several new molecular regulators of BBB function (Fig. 2), a level of speed and throughput not achievable with conventional tracer-based assays. In addition to chemical screening, FishNAP can assess BBB function in genetic mutants (Fig. 1). Because the assay is nonlethal and behavior-based, larvae displaying strong barrier phenotypes can be raised to establish stable lines with increased efficiency, making FishNAP well-suited for forward genetic screens that would be impractical in higher vertebrate models. Consistent with known BBB biology, FishNAP sensitively detects barrier changes across developmental and genetic contexts. By assaying larvae daily from 3–7 dpf, we found that the zebrafish BBB does not become functionally sealed until 5 dpf (Fig. 1B), as previously reported using classic tracer leakage assays^15^. FishNAP also robustly detects barrier defects in *spock1* mutants and *cldn5a* crispants, which model distinct disruptions in transcytosis regulation and tight junction organization (Fig. 1C & 1I)^8,19^. However, while we did see a significant difference between *cldn5a* and *EGFP* crispants in overall activity (Fig. 1I), we did not detect a significant correlation between *cldn5a* transcript levels and activity score (Fig. S2), likely reflecting noise inherent to both individual larval qPCR and single-fish FishNAP measurements. This suggests that FishNAP, in its current form, is best powered for group-level comparisons rather than fine-grained individual quantification, though it represents a promising starting point for further optimization toward single-animal resolution. Together, these results demonstrate that FishNAP reliably reports changes in barrier integrity across diverse biological contexts.

The compounds identified in this screen map to diverse signaling pathways—including Hedgehog, vitamin D, farnesyltransferase, cholesterol synthesis, receptor tyrosine kinases, and MEK–ERK signaling—highlighting that BBB integrity can be modulated through multiple regulatory pathways rather than a single canonical mechanism. Notably, multiple small molecules targeting these pathways were present in the screened library, yet only a subset altered BBB permeability (Fig. S5). This observation suggests that barrier modulation may depend on specific properties of individual compounds rather than pathway inhibition alone. For example, receptor tyrosine kinases comprise a large and diverse family, and it remains unclear whether the BBB-opening effects observed here reflect inhibition of particular signaling nodes, off-target activities, or structural features of the small molecules themselves. These findings therefore provide a starting point for future chemical and mechanistic studies aimed at defining the molecular determinants that enable selective modulation of BBB permeability. FishNAP can generate apparent false-positive hits because it relies on a behavioral readout of decreased locomotion. Although highly sensitive to neuroactive compounds, this readout is not specific for BBB permeability and may reflect off-target drug effects or drug interactions. This likely explains one of our validated hits, the P-glycoprotein (Pgp) inhibitor Berbamine HCl, which consistently sedated fish after 24 h of treatment (Fig. 2). Because Pgp actively exports many hydrophobic compounds from the brain^25,26^, its inhibition likely increased loperamide accumulation in the brain, producing sedation without widespread barrier leakage. Similarly, the apparent hits observed with the EGFR inhibitors Afatinib dimaleate and Poziotinib may reflect enhanced sedation rather than increased BBB permeability, as fatigue is a known side effect of these therapies^27,28^. These findings underscore the importance of validating FishNAP hits with tracer-based assays to determine whether permeability into the brain truly increases and whether such changes exhibit size or regional restrictions^15^. Because these compounds are delivered systemically, it is possible that other vascular interfaces are affected beyond the BBB. However, our analysis of peripheral tissues did not reveal any significant alterations in vascular permeability for either tracer (Fig. S4), suggesting that the effects are more selective for the brain. Future mechanistic studies using electron microscopy could help resolve more subtle vascular changes in peripheral tissues. Additionally, the brain is protected by multiple barriers, including the arachnoid barrier at the meninges and the blood–CSF barrier at the choroid plexus that may also be affected by these small molecules^29^. Neither FishNAP nor our tracer assays distinguish between these potential routes of entry into the brain. Defining which barriers are disrupted will require further mechanistic studies, including electron microscopy, long-term time-lapse imaging, transgenic permeability reporters, and single-cell transcriptomic profiling of different barrier-associated cells^8,15,17,30–32^.

Vitamin D signaling emerged as a potent regulator of BBB permeability in our screen, with direct vitamin D receptor agonists Calcifediol and Calcitriol reliably and reversibly increasing barrier permeability in zebrafish (Fig. 3 and 4), and Calcitriol also opening the mammalian barrier to the largest extent (Fig. 5). Additional members of the vitamin D metabolic pathway failed to alter BBB permeability via FishNAP (Fig. S6), which may be due to differences in effective dose in the brain as many of these compounds are metabolic precursors to Calcitriol. These findings are somewhat unexpected given prior studies linking vitamin D deficiency to BBB dysfunction and suggesting that Calcitriol treatment can stabilize barrier integrity under chronic deficiency conditions^33–37^. Our results point to a more complex relationship between vitamin D signaling and BBB regulation, in which acute increases in vitamin D activity can transiently increase barrier permeability. Together, these observations suggest that the magnitude, duration, or cellular context of vitamin D signaling may influence whether barrier integrity is reinforced or transiently relaxed. More broadly, the diverse pathways uncovered in our screen suggest that BBB permeability can be modulated through multiple regulatory inputs rather than a single dominant pathway. Critically, three compounds acting through distinct pathways, Calcitriol (vitamin D), Lovastatin (cholesterol synthesis), and Sunitinib (receptor tyrosine kinases), also increased BBB permeability in the adult mouse brain through both tight junction breakdown and increased transcytosis (Fig. 5 and 6), demonstrating that mechanisms uncovered through zebrafish screening translate directly to the mammalian BBB. This cross-species conservation underscores the power of FishNAP as a platform for *in vivo* drug discovery, capable of rapidly identifying BBB-modulating compounds in zebrafish that retain functional relevance in mammals, while also enabling the discovery and dissection of the signaling mechanisms that govern barrier permeability *in vivo*.

As in other areas of BBB research, findings from any single model are best interpreted in light of what that model can and cannot address. Zebrafish larvae share core features of BBB biology with mammals^1,15^, including a highly conserved barrier formed by endothelial cells supported by pericytes, but important differences exist. For example, astrocyte coverage of the vasculature is nearly complete in mammals but reaches only about 70% in zebrafish^38^. Physiology and drug delivery also differ: larvae rapidly absorb small molecules from the bath, whereas mammals are typically dosed by injection or other systemic routes, which makes direct comparison of effective doses and pharmacokinetics difficult. We therefore view FishNAP as a platform for rapid discovery that complements, rather than replaces, mammalian studies. Further mechanistic work will be needed to determine exactly how these molecules act, including whether they disrupt tight junctions, alter transcytosis, or both, and whether BBB opening is reversible and safe in mammalian models. Because FishNAP can rapidly narrow a large chemical space to a small set of candidates, it provides a strong foundation for these targeted studies, including efforts aimed at controlled, transient delivery of therapeutics to the brain.

## MATERIALS AND METHODS

### Zebrafish Lines and Maintenance

Zebrafish were maintained at 28.5°C following standard protocols^39^. All work was approved by the Rutgers University Standing Committee under protocol number 202300038. Adult fish were maintained on a standard 12 hour light-dark cycle. Adult fish, aged 3 months to 1.5 years, were crossed to produce embryos and larvae. For larvae used for confocal microscopy, 0.003% phenylthiourea (PTU; Sigma P7629) was applied beginning at 1 dpf to inhibit melanin production.

These studies used the AB wild-type strain, the transgenic reporter strains Tg(*kdrl:HRAS-mCherry*)^s896Tg^ abbreviated as *kdrl:mCherry* in the text^40^ and *Tg(l-fabp:DBP-EGFP)*^Iri500Tg^ abbreviated as *l-fabp:DBP-EGFP* in the text^41^, and the *spock1*^*hm43*^ BBB mutant line^8^. Crispant fish were generated by injection of Cas9 protein and multiple guide RNAs (IDT; *cldn5a* target sequences: GGACAACGTGAAAGCGCGGG, AGCGATGGCCTCCGCGGCTT, CAAGCACGAAGATGGCGCTG, GTGCGTCTGCGGGACGCTTT; *EGFP* target sequences: GGGCGAGGAGCTGTTCACCG and GAGCTGGACGGCGACGTAAA) into single-cell fertilized transgenic Tg(*kdrl:mCherry*) embryos in the AB background.

### FishNAP Assay

Fish were transferred to a 96 well square bottom plate, either individually or with 5-7 fish per well, and treated with 0.1 mM loperamide (Santa Cruz Biotechnology sc-203116) in 1x Danieau buffer immediately preceding the trial. The plate was then placed in a Daniovision recording chamber (Noldus). For grouped fish, mechanical tapping occurred every 10 seconds and lights flashed every 20 seconds. For individual fish, we used “rave” mode, with random tapping between 0-3 seconds and random light flashes within 0-10 seconds to encourage activity. Larval locomotor activity was recorded for 1 hour in 10-minute bins and quantified using Ethovision XT13 software (Noldus). An activity ratio was calculated by dividing the average activity during the final 20 minutes by the average activity during the first 20 minutes. Each condition was normalized to same-day controls (wild-type, uninjected, or DMSO-treated) to minimize batch effects, and normalized activity values were then tested for statistical differences.

### RNA isolation and RT-qPCR

The 96 crispant larvae were individually snap-frozen immediately following FishNAP. Total RNA was isolated using the Quick-RNA kit (Zymo Research) according to the manufacturer’s instructions. Individual cDNA was synthesized from 200 ng of total RNA using SuperScript IV Synthesis kit (Thermo Fisher Scientific), following the manufacturer’s protocol. 1.5 µl of cDNA was used per sample in a 6□µL reaction containing 0.5□µL of forward and reverse primers (5□mM each), 3□µL of SYBR® Green FastMix® (Quantabio) and run in 384-well format using a CFX Opus 384 DX instrument (Bio-Rad). PCR cycling profile consisted of a denaturing step at 95□°C for 1 min and 40 cycles at 95□°C for 10□s and 60□°C for 30□s. Ribosomal *18s* mRNA was used as normalization controls for 2^− ΔΔCt^ fold change analysis in comparison to *cldn5a* levels to determine relative levels of *cldn5a*. The sequence for the primers (IDT) used for *cldn5a* are *5’-* TGGAGCTCCTGGGTCTGATCCTGTG-3’ and 5’-GCACAGTGGCACAAGCACGAAGATG-3’ and for 18s are 5’-TCGCTAGTTGGCATCGTTTATG-3’ and 5’-CGGAGGTTCGAAGACGATCA-3’.

### Drug Screen

Wild-type zebrafish larvae were allocated into groups of seven per well in 24-well plates at 5 dpf. Larvae were exposed to 5 µM of two pooled compounds from the DiscoveryProbe™ FDA-approved Drug Library (APExBIO L1021) for two days and monitored for survival. At 7 dpf, larvae were analyzed using the FishNAP assay, with hits defined by a normalized activity ratio < 0.6. Pooled hits or toxic pools were retested with individual compounds at 5 µM to identify specific active molecules. Compounds exhibiting toxicity at 5 µM were then retested at 1 µM and subsequently at 0.5 µM with persisting toxicity. Hit validation was then performed in multiple wells across at least two independent experiments, with activity compared to DMSO-treated controls for significance.

### Zebrafish Fluorescent Tracer Leakage Assays and Image Analysis

Zebrafish larvae were anesthetized with tricaine and positioned in an agarose injection mold. Fish were injected with 2.3 nL containing Alexa 405 NHS Ester (10 mg/mL; Thermo Scientific A30000), Alexa 647 10 kDa Dextran (5 mg/mL; Thermo Scientific D22914), 70 kDa Dextran Fluorescein (5 mg/mL; Thermo Scientific D1822), or 376 Da fluorescein sodium (10 mg/mL; Sigma-Aldrich catalog #F6377-100G) into the cardiac sac using a Nanoject III microinjector (Drummond Scientific). Larvae were then embedded in 2% low-melting-point agarose (Promega V2111) and submerged in tricaine-containing embryo water for live imaging within 1 h post-injection. Imaging was performed on a Leica Stellaris DMi8 confocal microscope with a 25x water-immersion objective (0.75x zoom) and 1 µm z-steps. For BBB permeability assays, a 20 µm maximum-intensity projection starting ∼30 µm beneath the skin was used in FIJI^42^. Tracer intensity in the midbrain was quantified and normalized to intravascular tracer intensity for each fish for every tracer except the sodium fluorescein, which was normalized to the skin intensity. For trunk permeability analyses, a 60 µm maximum-intensity projection was made to encompass fin and axial muscle tissue in FIJI^42^. Tracer intensity in either the fin or muscle was quantified and normalized to intravascular tracer intensity for each fish. Representative images were made using 60 µm thick maximum-intensity projections.

### Mouse lines and manipulations

All mouse experiments were conducted in accordance with institutional and NIH guidelines and approved by the Rutgers University Standing Committee (protocol□#202400110). Mice were housed in a controlled indoor environment with a constant temperature of 22-23°C, humidity of 60%, and a 12□h light/12□h dark cycle. Wild-type C57BL/6J mice (Jackson Laboratory, strain 000664) aged 6–22 weeks, both male and female, were used. Calcitriol (1.2 µg g^−1^ body weight; Sigma-Aldrich 32222-06-3), Lovastatin (1.2 µg g^−1^ body weight; Aladdin Scientific L107709), or Sunitinib (15 µg g^−1^ body weight; SelleckChemicals 557795-19-4) was administered intraperitoneally once daily for two days. On day 3, mice received a retro-orbital injection of EZ-Link sulfo-NHS-LC-biotin tracer (0.2 mg g^−1^ body weight; Thermo Scientific 21335) under brief isoflurane anesthesia. Brains were collected 30 min later following rapid perfusion with 4% PFA, bisected and post-fixed overnight in either PFA or methanol at 4°C and subsequently processed for immunohistochemistry.

### Immunohistochemistry and Image Analysis

Dissected brain tissue was washed three times in PBS, equilibrated in 30% sucrose at 4°C, and flash-frozen in Tissue-Tek® O.C.T. compound (Sakura #4583) for cryosectioning at 20 μm thickness. Sagittal sections were blocked for 1 h at room temperature in PBS containing 10% normal donkey serum and 0.1% Triton X-100, then incubated overnight at 4°C with primary antibodies diluted in blocking buffer: goat anti-CD31 (1:100; R&D Systems AF3628), rabbit anti-CLDN5 (1:100; Invitrogen 34-1600), rabbit anti-MFSD2A (1:100; Cell Signaling Technologies 80302S), rabbit anti-CAV1 (1:100; Cell Signaling Technologies 3238S), rabbit anti-Albumin (1:50; abcam AB207327), and Donkey-anti-mouse IgG 488 (1:400 Jackson ImmunoResearch 715-545-151). Sections were subsequently washed in PBS and incubated for 2–3 h at room temperature with secondary antibodies (1:400; Jackson ImmunoResearch) and DyLight 405 Streptavidin (1:400; Jackson ImmunoResearch 016-470-084), then washed again in PBS and coverslipped.

Three cortical regions per mouse were imaged using a Leica Stellaris DMi8 confocal microscope with a 25× water objective (0.75x zoom). Maximum intensity projections were generated from 20 μm thick z-stacks and analyzed in FIJI for tracer permeability and vascular expression of CLDN5, MFSD2A and CAV1. Tracer permeability was calculated as the mean NHS, Albumin, or IgG intensity outside the vasculature normalized to that within CD31+ vessels, with values normalized to DMSO-treated controls from the same day (a value of 1 indicates no leakage; >1 indicates increased leakage). CLDN5+, MFSD2A+, and CAV1+ vasculature was quantified by thresholding CD31+ regions to define vascular ROIs, then measuring the % area containing Huang-thresholded CLDN5, MFSD2A, or CAV1 signal.

### Statistical Analysis

All statistical analyses were performed using Prism 10 (GraphPad Software). Appropriate statistical tests were determined according to the number of comparisons, the number of variables and the appropriateness of normality assumptions. Sample size for all experiments was determined empirically using standards generally employed by the field, and no data was excluded when performing statistical analysis. ∗∗∗∗p < 0.0001, ∗∗∗p < 0.001, ∗∗p < 0.01, ∗p < 0.05 as stated in the Figure Legends. Standard error of the mean was calculated for all experiments and displayed as errors bars in graphs. Statistical details for specific experiments, including precision measures, statistical tests used, and definitions of significance can be found in the figure legends.

## ACKNOWLEDGEMENTS

We thank members of the O’Brown lab for data discussions and comments on the manuscript; Dr. Zach O’Brown for discussions and comments on the manuscript; and the Aquatic Core facility, specifically Katie Flaherty, at Rutgers for fish maintenance and care. We also thank Drs. Rebecca Ward and Randall Peterson for initial brainstorming and discussions that enabled FishNAP to become a reality.

## AUTHOR CONTRIBUTIONS

N.M.O. and T.C.P. developed the concept, experimental design, and wrote the manuscript. C.G.R. contributed to the development of FishNAP. T.C.P., E.E.M., J.L., L.G.G., E.C., N.G., A.K.V., A.R.I., M.H.G., K.A.P., and M.E.C. conducted animal experiments and analyzed data.

## DATA AND CODE AVAILABILITY

The datasets generated and analyzed during this study are provided within the manuscript and its supplementary information, including the complete FishNAP screening dataset (Supplementary Data 1). Owing to their size (several hundred gigabytes), the raw imaging data, comprising full z-stacks for each compound and control, are available from the corresponding author upon request. Source data are provided with this paper.

