## Supplementary Information for "In vivo discovery of blood-brain barrier opening small molecules with FishNAP"

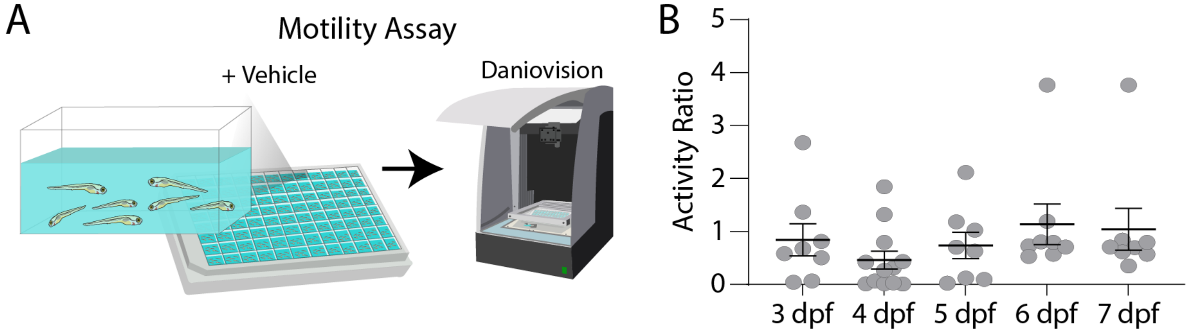


**Fig. S1. Larval zebrafish show no change in baseline activity. A.** Schematic of baseline motility assay using the same FishNAP settings in the absence of loperamide. **B.** Quantification of activity across larval development reveals no change in baseline activity at any stage. Each data point represents a well of 7 fish. Bars show the mean ± SEM. No significance by Kruskal-Wallis test.

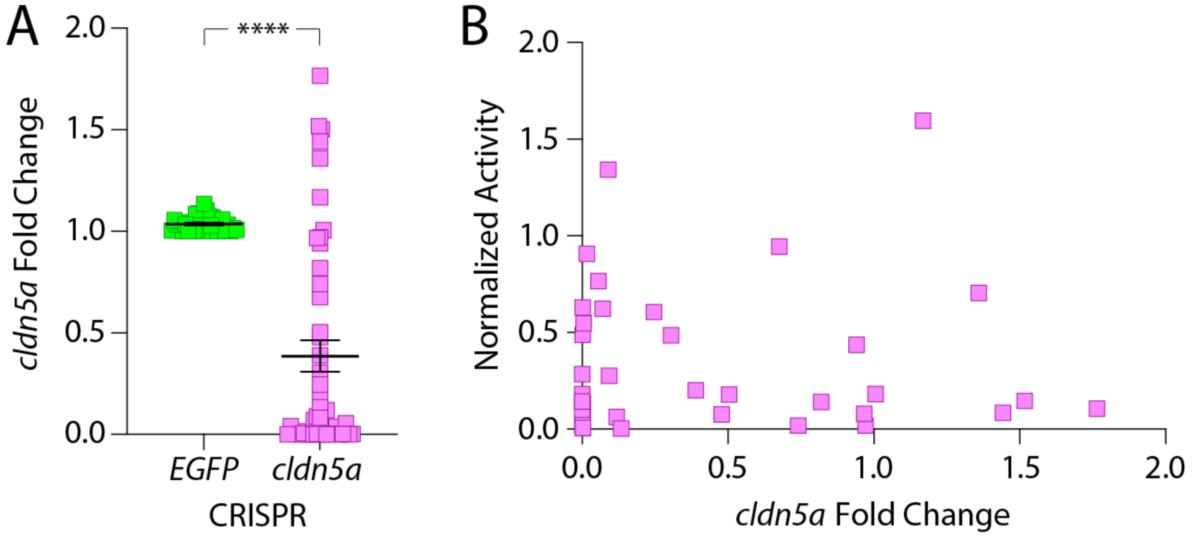


**Fig. S2. *Cldn5a* crispants have a significant reduction in *cldn5a* expression. A.** Compared to *EGFP* crispant controls, *cldn5a* crispants have a significant reduction in *cldn5a* levels. **B.** There is no significant correlation between *cldn5a* levels and normalized activity from FishNAP. Each square data point represents an individual fish. Bars show the mean ± SEM. **** $p<0.0001$ by Welch's t-test.

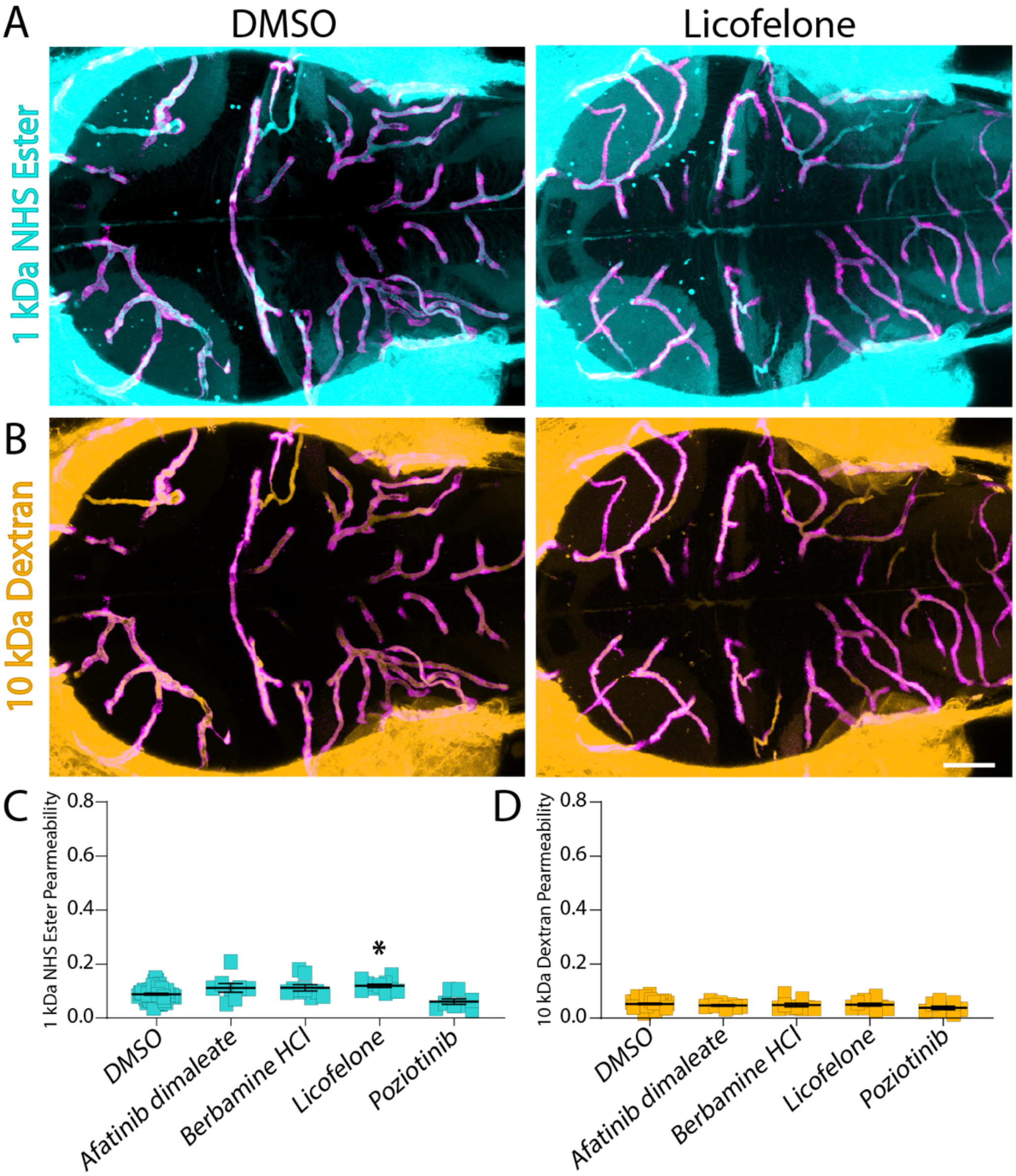


**Fig. S3. Four small molecules fail to cause substantial leakage of the BBB. A-B.** Functional tracer leakage assays with 1 kDa NHS Ester (turquoise, A) and 10 kDa Dextran (gold, B) reveal a small increase in 1 kDa NHS leakage after Licofelone treatment. **C-D.** Quantification of leakage in all treated fish for 1 kDa NHS Ester (C) and 10 kDa Dextran (D). Each point represents an individual fish. Bars show the mean ± SEM. * $p<0.05$ by one-way ANOVA with Dunn’s post-hoc test compared to DMSO controls. Scale bar represents 50 µm.

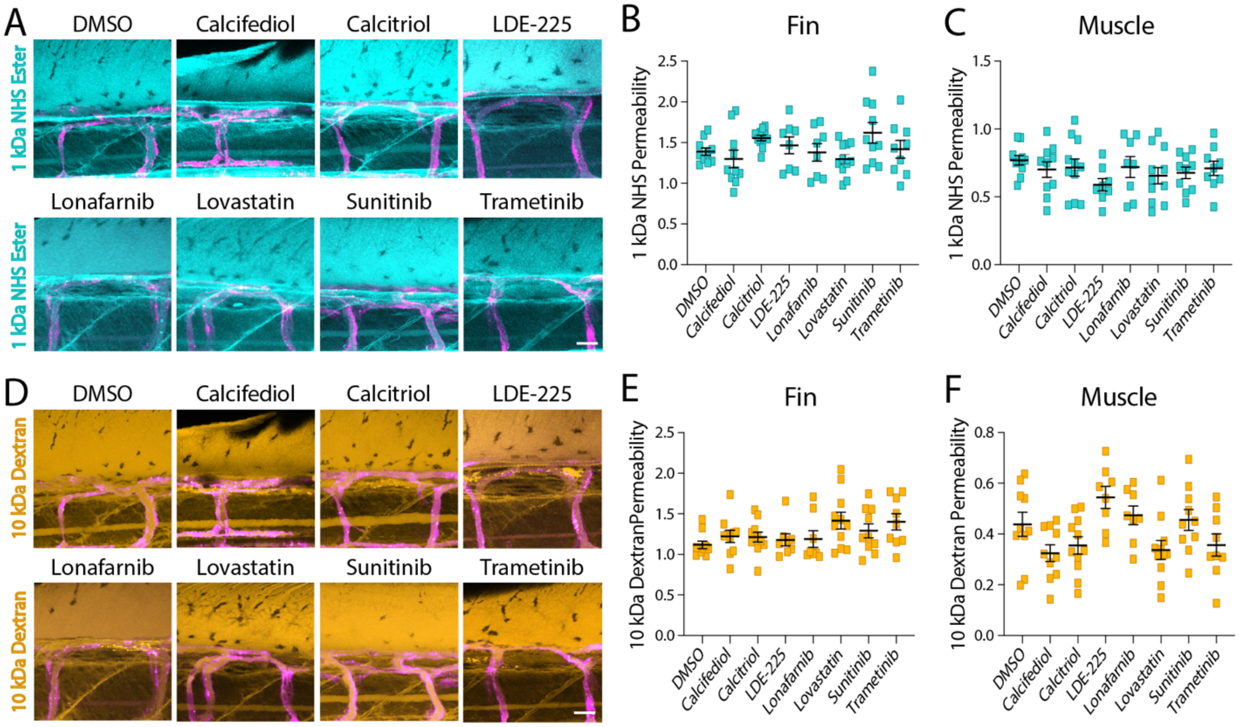


**Fig. S4. Small molecule treatment does not alter vascular permeability in peripheral tissues. A-C.** Larvae were treated with each small molecule from 5–7 dpf and assayed for leakage of 1 kDa NHS Ester (turquoise) from the vasculature (magenta) into the fin (B) and muscle (C) tissue, revealing no change in permeability compared to DMSO controls. **D–F**. Larvae were treated with each small molecule from 5–7 dpf and assayed for leakage of 10 kDa Dextran (gold, D) from the vasculature (magenta) into the fin (E) and muscle (F) tissue, revealing no change in permeability compared to DMSO controls. Each square data point represents an individual fish in B, C, E, and F with a minimum of 8 larvae tested per condition. Bars show the mean ± SEM. No significance by one-way ANOVA. Scale bars represent 20 μm.

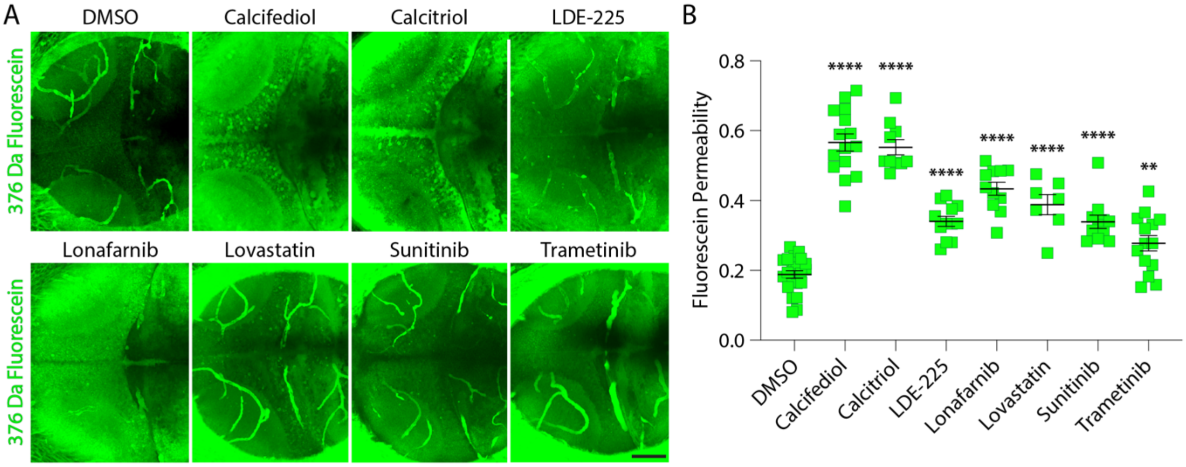


**Fig. S5. All seven BBB-opening compounds increase permeability to a 376 Da Fluorescein tracer. A.** Larvae were treated with each small molecule from 5–7 dpf and assayed for leakage of 376 Da Fluorescein Sodium (green) into the brain parenchyma, revealing increased permeability compared to DMSO controls. **B.** Quantification of Fluorescein permeability reveals that all seven molecules significantly increase permeability to the small 376 Da tracer, similar to the 1 kDa NHS Ester tracer. Each square data point represents an individual fish, with a minimum sample size of 7 larvae. Bars show the mean ± SEM. ** $p<0.01$, **** $p<0.0001$ by one-way ANOVA with Dunn's post-hoc test compared to DMSO controls. Scale bar represents 50 µm.

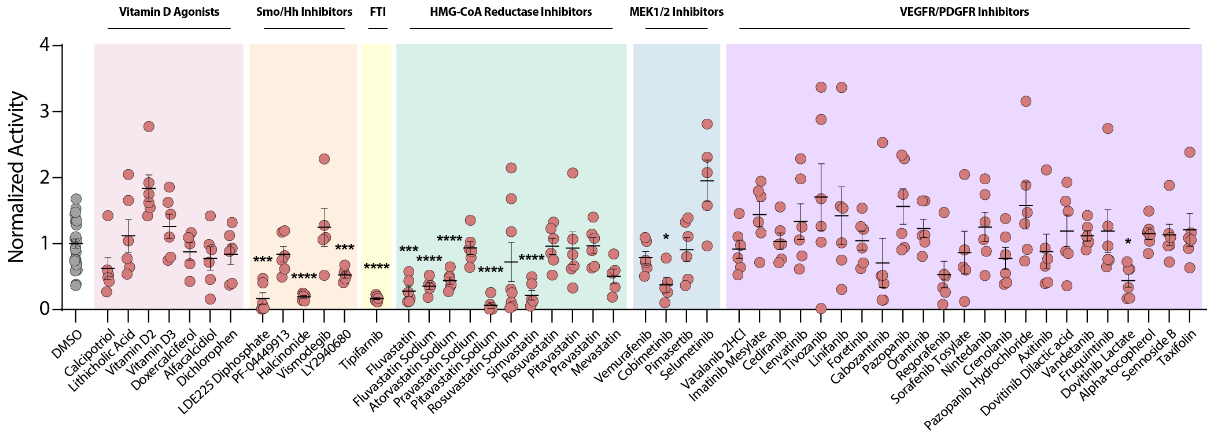


**Fig. S6. FishNAP screening of related signaling pathway members identifies additional BBB-opening candidates.** Further testing of other vitamin D agonists revealed no further candidates beyond Calcitriol and Calcifediol. Three additional smoothend (Smo)/hedgehog (Hh) inhibitors were identified: Halcinonide, LDE225 Diphosphate and LY2940680. The only other farnesyltransferase inhibitor (FTI) tested, Tipifarnib, significantly increased sedation by FishNAP, similar to Lonafarnib. One other MEK1/2 inhibitor in addition to Trametinib, Cobimetinib, had moderate, but significant effects on sedation. Several other lipophilic HMG-CoA reductase inhibitor candidates were identified, including Fluvastatin, Pitavastatin Calcium, and Simvastatin, while no hydrophilic statins had an effect. Only one other VEGFR/PDGFR inhibitor in addition to Sunitinib had an effect: Dovitinib Lactate. Each data point represents a well of 7 treated fish with a minimum of 5 wells (35 larvae) tested per treatment. Bars show the mean ± SEM. * $p<0.05$, *** $p<0.001$, **** $p<0.0001$ by Welch's ANOVA with Dunnett's T3 post-hoc test compared to DMSO controls.

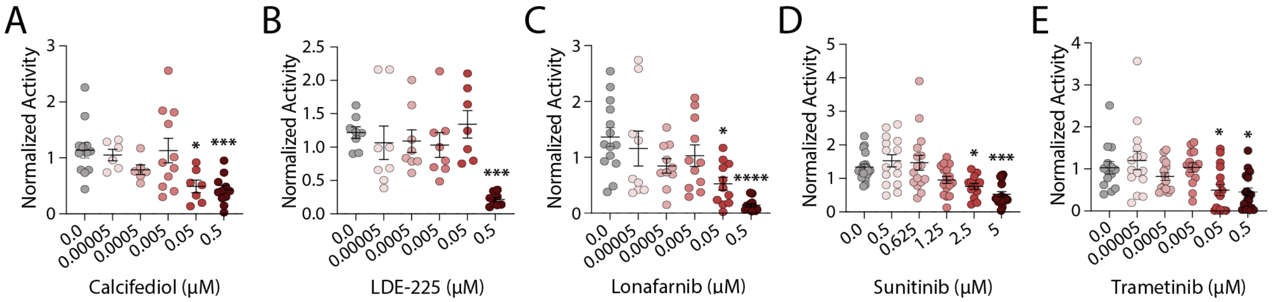


**Fig. S7. Identifying the minimum effective doses that reliably increase BBB permeability for each small molecule. A**. Calcifediol is effective at both 0.05 μM and 0.5 μM. **B.** LDE-225 is effective only at the screening concentration of 0.5 μM. **C.** Lonafarnib is effective at both 0.05 μM and 0.5 μM. **D.** Sunitinib is effective at both 2.5 μM and 5 μM. **E.** Trametinib is weakly effective at both 0.05 μM and 0.5 μM. Each data point represents a well of 7 fish, with a minimum of 7 wells (49 larvae) tested per condition. Bars show the mean ± SEM. * $p<0.05$, *** $p<0.001$, **** $p<0.0001$ by Kruskal-Wallis test with Dunn's post-hoc test compared to DMSO controls.
